# Phosphorylated tau fluid biomarker sites recognize earlier neurofibrillary tangle maturity levels in the postmortem Alzheimer’s disease brain

**DOI:** 10.1101/2021.08.25.457363

**Authors:** Christina M. Moloney, Sydney A. Labuzan, Julia E. Crook, Habeeba Siddiqui, Monica Castanedes-Casey, Christian Lachner, Ronald C. Petersen, Ranjan Duara, Neill R. Graff-Radford, Dennis W. Dickson, Michelle M. Mielke, Melissa E. Murray

## Abstract

Alzheimer’s disease (AD) biomarkers have become increasingly more reliable in predicting AD pathology. While phosphorylated tau fluid biomarkers have been studied for over 20 years, there is a lack of deep characterization of these sites in the postmortem brain. Neurofibrillary tangle-bearing neurons, one of the major neuropathologic hallmarks of AD, undergo morphologic changes that mature along a continuum as hyperphosphorylated tau aggregates. To facilitate interpretation of phosphorylated tau sites as an early fluid biomarker, our goal was to characterize which neurofibrillary tangle maturity levels (pretangle, intermediary 1, mature tangle, intermediary 2, and ghost tangle) they recognize. We queried the Florida Autopsied Multi-Ethnic (FLAME) cohort for cases from Braak stages I-VI. We excluded non-AD pathologies and tauopathies. A total of 24 cases, 2 males and 2 females for each Braak stage, were selected. We performed immunohistochemistry on the posterior hippocampus using antibodies directed towards phospho (p) threonine (T) 181, pT205, pT217, and pT231. Slides were digitized to enable quantification of tau burden. To examine differences in regional vulnerability between CA1 and subiculum, we developed a semi-quantitative system to rank the frequency of each neurofibrillary tangle maturity level. We identified all neurofibrillary tangle maturity levels at least once for each phosphorylated tau site. Primarily earlier neurofibrillary tangle maturity levels (pretangle, intermediary 1, mature tangle) were recognized for all phosphorylated tau sites. There was an increase in tau burden in the subiculum compared to CA1; however, this was attenuated compared to thioflavin-S positive tangle counts. On a global scale, tau burden generally increased with each Braak stage. These results provide neurobiologic evidence that these phosphorylated tau fluid biomarker sites are present during earlier neurofibrillary tangle maturity levels. This may help explain why these phosphorylated tau biomarker sites are observed before symptom onset in fluids.

## Introduction

Alzheimer’s disease (AD) is a neurodegenerative disorder characterized by two hallmark pathologies: plaques composed of amyloid-β and neurofibrillary tangles composed of hyperphosphorylated tau [1, 2, 49]. Amyloid-β plaques reside outside the neuron whereas neurofibrillary tangles accumulate inside the neuron. Neurofibrillary tangles have a lifespan that progresses through three major levels: pretangles, mature tangles, and ghost tangles [42]. Pretangles contain diffuse or granular hyperphosphorylated tau in otherwise morphologically normal neurons with perinuclear tau accumulation often observed [6]. Mature tangles are intensely stained bundles of fibers that take the shape of the neuron they occupy. The nucleus may appear shrunken and misplaced towards the cell membrane [1, 2, 49]. Ghost tangles are the remnants of mature tangles once the neurons have died and are recognized by faintly stained bundles of fibers with no associated nucleus [1, 2, 49]. Additionally, intermediaries exist between these three neurofibrillary tangle maturity levels [42]. The first intermediary, between pretangles and mature tangles, has intensely stained tau accumulation in the neuron, but lacks the fullness of the mature tangle. These were previously described as nucleation sites [33] that may represent a nidus of fibrillization. The second intermediary, between mature tangles and ghost tangles, appears robustly stained like mature tangles but lacks a nucleus.

Biomarkers are an important tool that have become increasingly more reliable in measuring AD pathology in vivo [20]. One such biomarker is positron emission tomography (PET), which can identify patients with increased tau or amyloid-β levels. PET imaging can lead to a more informed antemortem diagnosis and provide spatial information of which regions have a high neuropathologic burden [31, 44]. However, disadvantages to PET imaging include cost, availability, and patient exposure, albeit minimal, to radioactivity. Alternatively, fluid biomarkers are useful in identifying changes in amyloid-β and tau in patients without the use of radioactivity [40]; however, these biomarkers lack spatial information. Over 20 years ago, total tau [38, 56, 59] and phosphorylated tau [11] were found increased in cerebrospinal fluid of AD patients compared to controls. Multiple tau phosphorylation (p) sites are elevated in these fluid biomarkers, including at threonine (T) 181 [11, 57], pT205 [9], pT217 [7, 39], and pT231 [11, 24]. Evaluation of the temporal sequence of cerebrospinal fluid tau biomarkers in dominantly inherited AD identified pT217 as increasing first, followed by pT181, and finally pT205 [8]. Following the successful translation of cerebrospinal fluid assays to plasma [4, 41, 45, 52], one study suggested pT231 levels may increase earlier than pT181 [4]. With the increasing evidence that tau fluid biomarkers may be more sensitive to the earlier changes in AD pathology, we sought to test the hypothesis that these phosphorylation sites recognize earlier neurofibrillary tangle maturity levels. Thus, our first goal was to immunohistochemically characterize the neurofibrillary tangle maturity levels recognized by these four fluid biomarker phosphorylated tau sites: pT181, pT205, pT217, and pT231. Secondly, we sought to examine regional differences leveraging the vulnerability of cornu ammonis (CA) 1 and subiculum [43, 46] using digital pathology to quantify tau burden. Lastly, we sought to identify changes in the phosphorylated tau sites based on global severity measured by Braak stage [12, 13] to evaluate shifting recognition of neurofibrillary tangle maturity levels.

## Materials and Methods

### Cohort selection

The Florida Autopsied Multi-Ethnic (FLAME) cohort [28, 47], as of May 27, 2020, was queried to identify cases with a spectrum of AD neuropathology. Cases with significant non-AD neurodegenerative pathology (frontotemporal lobar degeneration, amyotrophic lateral sclerosis, hippocampal sclerosis, Lewy body disease, amygdala predominant Lewy bodies, multiple system atrophy, Creutzfeldt-Jakob disease) and non-AD tauopathies (progressive supranuclear palsy, corticobasal degeneration, Pick’s disease, globular glial tauopathies, *MAPT* mutation carriers) were excluded. Cases with exhausted paraffin embedded blocks of the posterior hippocampus were also excluded. Ethnoracial minorities were given higher priority when selecting cases as these groups have historically been excluded from studies. We selected 2 males and 2 females for each Braak stage I, II, III, IV, V, and VI [12, 13]. For Braak stages IV-VI, only AD cases with a typical AD subtype were selected [43]. A summary of cases included in this study is found in **Supplementary Table 1**. The final cohort included 24 cases (**Table 1**). There were 2 males and 2 females per Braak stage. There was 1 black decedent (Braak stage III), and 5 Hispanic decedents (1 Braak stage I, 1 Braak stage III, 2 Braak stage V, 1 Braak stage VI). There were 7 *APOE* ε4 positive cases (2 Braak stage IV, 3 Braak stage V, 2 Braak stage VI).

**Table 1.** Characteristics and digital pathology findings by Braak stage.

| Characteristic | Braak I | Braak II | Braak III | Braak IV | Braak V | Braak VI | p-value |
| --- | --- | --- | --- | --- | --- | --- | --- |
| Number | 4 | 4 | 4 | 4 | 4 | 4 |  |
| Age at death, year | 87 (79, 93) | 80 (71, 84) | 89 (87, 93) | 84 (82, 93) | 86 (82, 90) | 79 (71, 84) | 0.041 |
| Females (%) | 2/4 (50%) | 2/4 (50%) | 2/4 (50%) | 2/4 (50%) | 2/4 (50%) | 2/4 (50%) | 1.000 |
| *APOE* ε4+ (%) | 0/4 (0%) | 0/4 (0%) | 0/4 (0%) | 2/4 (50%) | 3/4 (75%) | 2/4 (50%) | 0.058 |
| Brain weight, g | 1305 (960, 1440) | 1090 (1060, 1220) | 1140 (1000, 1320) | 1050 (1020, 1340) | 1030 (960, 1160) | 890 (760, 1100) | 0.002 |
| **Digital pathology findings** |  |  |  |  |  |  |  |
| Whole hippocampus measures |  |  |  |  |  |  |  |
| Area (mm<sup>2</sup>) | 44 (43, 49) | 45 (39, 60) | 50 (35, 67) | 48 (35, 67) | 31 (28, 54) | 34 (31, 40) | 0.034 |
| Positive Pixel Count |  |  |  |  |  |  |  |
| pT181 burden, % | 1.6 (0.50, 7.9) | 2.9 (0.47, 7.7) | 1.7 (0.64, 4.5) | 4.1 (2.8, 8.6) | 7.2 (1.8, 8.9) | 13 (10, 14) | <0.001 |
| pT205 burden, % | 0.52 (0.15, 0.82) | 0.19 (0.065, 4.7) | 2.3 (1.2, 5.8) | 10 (2.7, 13) | 9.4 (6.7, 12) | 9.5 (6.8, 12) | <0.001 |
| pT217 burden, % | 0.78 (0.38, 2.0) | 0.70 (0.090, 7.4) | 4.7 (2.9, 10) | 16 (5.6, 18) | 13 (12, 17) | 20 (8.3, 23) | <0.001 |
| pT231 burden, % | 0.25 (0.17, 0.36) | 0.11 (0.048, 2.3) | 1.5 (0.93, 3.5) | 6.6 (2.9, 8.1) | 8.3 (4.9, 13) | 12 (6.4, 17) | <0.001 |
| CA1 measures |  |  |  |  |  |  |  |
| Thioflavin S counts | 0 (0, 1) | 1 (0, 1) | 2 (1, 4) | 5 (2, 10) | 10 (4, 25) | 22 (6, 30) | <0.001 |
| Positive Pixel Count |  |  |  |  |  |  |  |
| pT181 burden, % | 0.95 (0.24, 3.5) | 1.4 (0.19, 3.6) | 1.8 (0.58, 3.2) | 3.8 (3.5, 12) | 7.3 (2.4, 11) | 14 (5.8, 18) | <0.001 |
| pT205 burden, % | 0.70 (0.25, 2.5) | 0.21 (0.066, 6.0) | 3.1 (1.7, 8.8) | 13 (3.8, 20) | 15 (11, 17) | 13 (8.9, 18) | <0.001 |
| pT217 burden, % | 0.78 (0.60, 3.0) | 0.37 (0.073, 9.6) | 6.2 (3.9, 17) | 25 (7.1, 28) | 19 (17, 25) | 30 (11, 33) | <0.001 |
| pT231 burden, % | 0.27 (0.24, 0.70) | 0.11 (0.046, 3.1) | 2.2 (1.4, 6.5) | 10 (3.6, 15) | 14 (6.4, 21) | 17 (7.9, 28) | <0.001 |
| Subiculum measures |  |  |  |  |  |  |  |
| Thioflavin S counts | 0 (0, 1) | 0 (0, 1) | 2 (1, 4) | 8 (6, 17) | 22 (8, 40) | 40 (25, 55) | < 0.001 |
| Positive Pixel Count |  |  |  |  |  |  |  |
| pT181 burden, % | 1.4 (0.37, 7.4) | 2.8 (0.26, 4.7) | 2.7 (0.91, 4.4) | 6.7 (4.6, 13) | 9.0 (3.5, 11) | 21 (13, 23) | <0.001 |
| pT205 burden, % | 1.1 (0.24, 1.2) | 0.24 (0.064, 9.9) | 4.9 (3.1, 12) | 19 (6.8, 20) | 18 (11, 21) | 18 (11, 22) | <0.001 |
| pT217 burden, % | 1.5 (0.64, 2.4) | 0.64 (0.082, 16) | 10 (6.9, 20) | 31 (13, 39) | 23 (17, 34) | 38 (16, 43) | <0.001 |

### Tissue sampling and histology

Autopsied brains were evaluated and sampled using the <u>Dickson</u> sampling scheme, as previously described [47]. Formalin-fixed, paraffin-embedded tissue sections of the posterior hippocampus were cut at 5 µm thick and mounted on positively charged glass slides. Thioflavin-S was used to assess AD pathology to assign AD subtype [43], Braak stage [13], and Thal phase [53]. Primary age related tauopathy (PART) was diagnosed if cases were Braak stage ≤IV with a “definite” classification if Thal phase 0 or a “possible” classification if Thal phase 1-2 [14]. Cases not meeting recommended PART criteria who were Thal phase >2 were diagnosed as pathological aging if significant cortical amyloid-β pathology (≥30 amyloid-β plaques per 10x field) with minimal neurofibrillary tangle pathology (Braak stage <IV) was observed [15]. Further, these Braak <IV cases not meeting recommended PART criteria who were Thal phase >2 with non-significant cortical amyloid-β pathology (<30 amyloid-β plaques per 10x field) were diagnosed as senile change [31]. Aging-related tau astrogliopathy (ARTAG) was diagnosed if cases had thorn-shaped or granular/fuzzy astrocytes [25]. Significant vascular disease was diagnosed in the presence of two or more of the following pathologies: severe cerebral amyloid angiopathy, moderate-to-severe white matter rarefaction, hippocampus sclerosis of a vascular etiology, large infarct, lacunar infarct, microinfarct, or white matter infarct.

Sections were stained with hematoxylin and eosin (H&E) using standard methods. For neurofibrillary tangle counts, sections were stained with thioflavin-S. Neurofibrillary tangles (mature tangles and ghost tangles) were collectively counted in a 40x field of view (0.125 mm^2^) with highest burden in the CA1 subsector of the hippocampus and the subiculum using the Olympus BH2 fluorescence microscope. Immunohistochemistry using AT270 (pT181), pT205, pT217, and AT180 (pT231) antibodies was performed on the Thermo Scientific Lab Vision Autostainer 480S. Specifications and vendor information for each antibody is found in **Table 2**. All epitopes fall within the proline rich region (**Fig. 1**), sometimes described as mid-domain, which lies N-terminal to the microtubule binding region [19]. Sections were developed with the developing kit (Biocare Medical, catalog #M3M530L [mouse] or M3R531L [rabbit]). Regarding nomenclature, the current study will use “pT” when describing immunohistochemical evidence from the phosphorylated threonine site and “p-tau” when discussing relevant information from the fluid biomarker literature on phosphorylated tau.

**Table 2.** Antibody information

| Antibody | Epitope | Vendor | Catalog # | Species | Clonality | Isotype | RRID | Dilution | Antigen Retrieval | Ref |
| --- | --- | --- | --- | --- | --- | --- | --- | --- | --- | --- |
| AT270 | pT181 | Thermo Fisher Scientific | MN1050 | Mouse | Monoclonal | IgG1κ | AB_223651 | 1:10,00 | H<sub>2</sub>O steam | [17] |
| pT205 | pT205 | Thermo Fisher Scientific | 44-738G | Rabbit | Polyclonal | IgG | AB_2533738 | 1:2,500 | H<sub>2</sub>O steam |  |
| pT217 | pT217 | Thermo Fisher Scientific | 44-744 | Rabbit | Polyclonal | IgG | AB_2533741 | 1:500 | H<sub>2</sub>O steam |  |
| AT180 | pT231 | Thermo Fisher Scientific | MN1040 | Mouse | Monoclonal | IgG1κ | AB_223649 | 1:5,000 | H<sub>2</sub>O steam | [17] |

**Figure 1.**
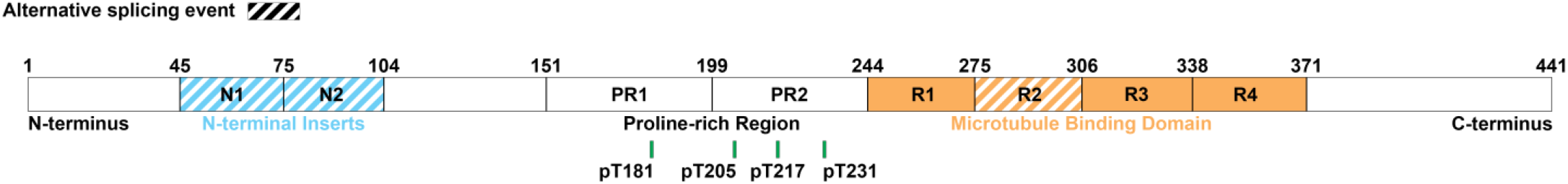
All phosphorylated tau antibodies used in this study fall within the proline-rich region. Amino acid position is illustrated by a green line for phosphorylated threonine (pT) at 181, 205, 217, and 231. Figure adapted from [42].

### Digital pathology

H&E and immunohistochemically stained slides were digitally scanned at 20x using the Aperio AT2 scanner. Using ImageScope (Leica Biosystems, version 12.4.3.5008), digitally scanned slides were annotated with the pen tool to trace the CA1 and subiculum. The superior border was defined as the boundary between the lacunosum and radiatum layers. The inferior border was defined as the boundary between the pyramidal layer and alveus. The border between CA1 and subiculum was defined as halfway between the dentate gyrus length. For consistency using landmarks, the CA1 was traced from the CA1/subiculum border to the ventricle. The whole hippocampus was annotated, including the alveus, excluding the fornix, and allowing the rise of the subiculum into the presubiculum to come to a natural endpoint. Hippocampal area (per mm^2^) was traced on each of the immunohistochemically stained sections and averaged to obtain a final area. H&E stained sections facilitate recognition of neuroanatomic boundaries and are thus traced first as a template for subsequent tracing on immunohistochemical slides. For all scans, dust and artifacts were excluded using the negative pen tool.

Batch analysis of the scans was completed using eSlideManager (Leica Biosystems). For burden analysis, digitized slides were analyzed with color deconvolution (v9) (**Supplementary Table 2**) and positive pixel count (v9) (**Supplementary Table 3**) macros custom designed for each antibody to recognize the 3, 3’-diaminobenzidine staining on the tissue and to exclude background. Positive pixel count results will be described in this study; for color deconvolution results, please see **Supplementary Table 4**.

### Neurofibrillary tangle maturity level classifications

Neurofibrillary tangles were classified into the following maturity levels: pretangle, intermediary 1 tangle, mature tangle, intermediary 2 tangle, and ghost tangle. Pretangles were identified by diffuse or granular staining of phosphorylated tau and could have perinuclear accumulation of tau. Pretangles were less intensely stained tau than mature tangles. Intermediary 1 tangles were identified with intensely stained aggregates that did not fill the entire neuron. These aggregates may be round or fibrillar in appearance. Mature tangles were identified by intense staining of phosphorylated tau throughout most, if not all the neuron, with an associated nucleus that was often displaced or shrunken in appearance. Intermediary 2 tangles were identified with intense staining of phosphorylated tau throughout the entire neuron but lack a nucleus. Ghost tangles were identified with weaker staining of phosphorylated tau compared to mature tangles in structures that appear to be loosely arranged bundles of fibers with no associated nucleus.

### Neurofibrillary tangle maturity semi-quantification

Neurofibrillary tangle maturity levels were semi-quantified in the annotated regions in the CA1 and subiculum using a grading system: absence, rare, and presence. Absence referred to no pathology that resembled a maturity level. Rare referred to 1-5 individual tangles of a certain maturity level. Presence referred to greater than 5 tangles of a certain maturity level. For quantification, we assigned each frequency a value: absence=0, rare=0.5, presence=1. Values from cases in each Braak stage per phosphorylated tau site were added together.

### Statistical analysis

To compare the four tau biomarker sites to each other, we analyzed all scans with both positive pixel count and color deconvolution (**Supplemental Fig. 1**). Continuous values are reported as median (interquartile range [25^th^, 75^th^]). A permutation test with Kendall’s Tau was used to test differences of continuous values between Braak stages. Spearman correlation was used for the whole hippocampal area measures with tau burden. A Pearson’s chi-squared test was used to test differences of categorical values between Braak stages. P-values <0.05 were considered statistically significant. For comparing CA1 and subiculum burden, a simple linear regression was performed using GraphPad Prism and ratio was assessed by slope of the best fit line (version 9.0.0). All other statistical analyses were performed using R (version 4.0.3).

### Figure development

Figures were created using Adobe Photoshop CC 2018 (version 19.1.6). Snapshots of 10x and 20x immunohistochemically stained sections were acquired using ImageScope (Leica Biosystems, version 12.4.3.5008). Thioflavin-S stained slides were digitally scanned using the Pannoramic 250 Flash III scanner (3DHistech) and snapshots of a 20x view of the CA1 and subiculum were taken using the CaseViewer software (3DHistech, version 2.3.0.99276) after digital alteration to increase the brightness for illustration purposes. The heatmap was created in Microsoft Excel (Microsoft 365, version 16.0.13127.21336). Graphs were created using GraphPad Prism (version 9.0.0).

## Results

### Morphologic characteristics of neurofibrillary tangle and neuritic pathology

To evaluate the neurofibrillary tangle maturity level for each phosphorylated tau site, we performed immunohistochemistry on serial sections of the hippocampus for each antibody. At least one instance of each neurofibrillary tangle maturity level was visualized for all phosphorylated tau sites (**Fig. 2**). In addition, we found neuritic pathology (neuropil threads, neuritic plaques, and tangle associated neuritic clusters) was recognized by all phosphorylated tau antibodies (**Fig. 2**). Pathologic features of ARTAG (thorn shaped astrocytes) and argyrophilic grains disease (coiled bodies) were also observed (**Supplementary Fig. 2**). To determine the predominance of neurofibrillary tangle maturity level, we semi-quantified the presence or absence of pretangles, intermediary 1s, mature tangles, intermediary 2s, and ghost tangles (**Fig. 3**). There was a predilection towards earlier neurofibrillary tangle pathology especially in earlier Braak stages. Ghost tangles were extremely rare for all phosphorylated tau sites.

**Figure 2.**
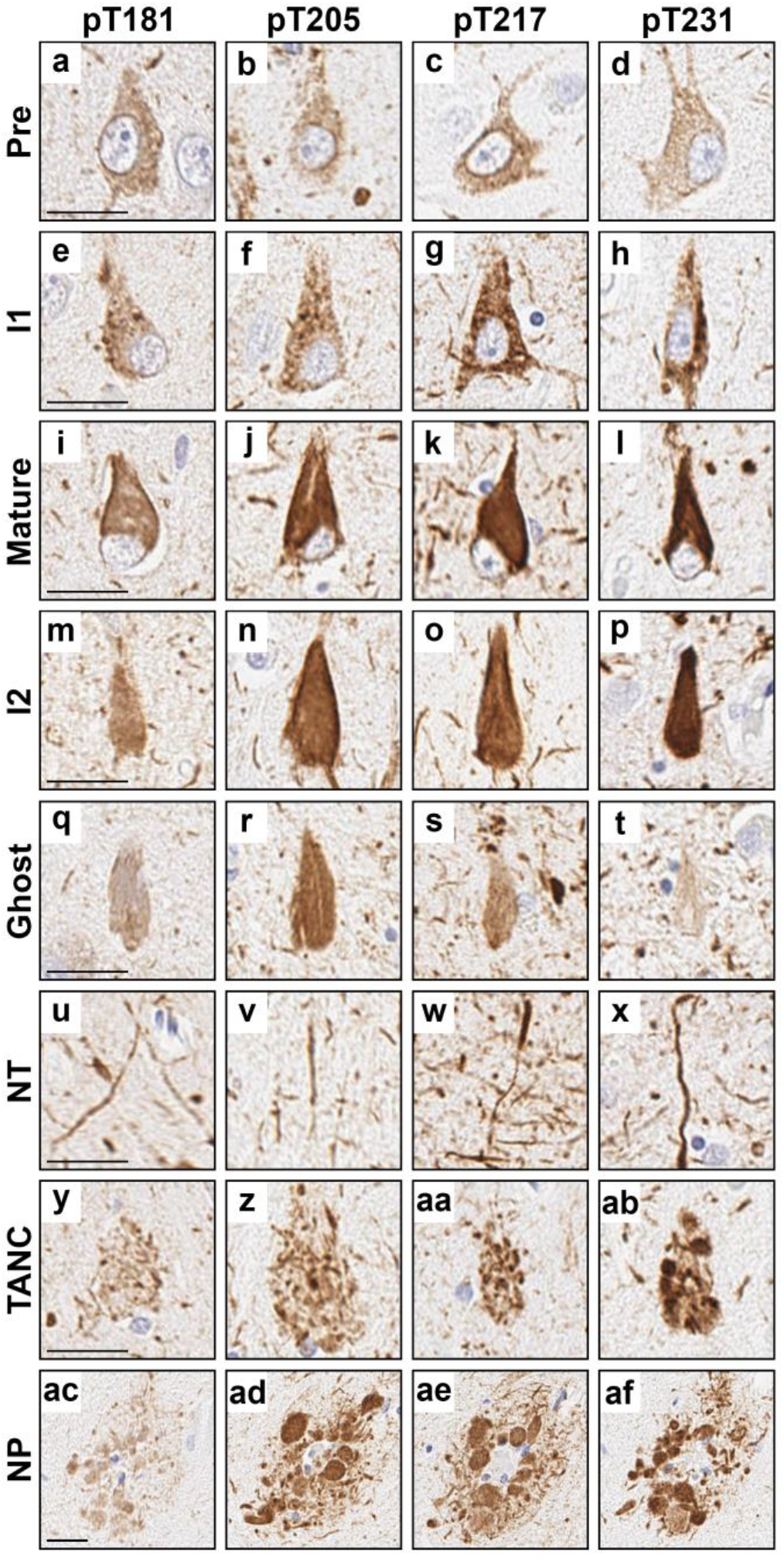
Examples of neurofibrillary tangle maturity levels and neuritic pathology observed with each phosphorylated tau site. **a-d** Pretangles, **e-h** intermediary 1 (I1), **i-l** mature tangles, **m-p** intermediary 2 (I2), **q-t** ghost tangles, **u-x** neuropil threads (NT), **y-ab** tangle associated neuritic clusters (TANC), and **ac-af** neuritic plaques (NP) were observed for all phosphorylated tau sites. **a, e, i, m, q, u, y**, and **ac** were stained with the pT181 antibody. **b, f, j, n, r, v, z**, and **ad** were stained with the pT205 antibody. **c, g, k, o, s, w, aa**, and **ae** were stained with the pT217 antibody. **d, h, l, p, t, x, ab**, and **af** were stained with the pT231 antibody. All images were taken in the CA1 subsector of the hippocampus except for **q, r**, and **t** which were taken in the subiculum. **a, e, i, l, t, k**, and **ab** are from case 18. **b** is from case 9. **c** is from case 14. **d, h, n**, and **ac-af** are from case 17. **f, j, v**, and **z** are from case 16. **g, q, r**, and **aa** are from case 13. **k, s, u, w**, and **y** are from case 15. **m, p**, and **x** are from case 23. **o** is from case 22. Scale bar measures 25 µm.

**Figure 3.**
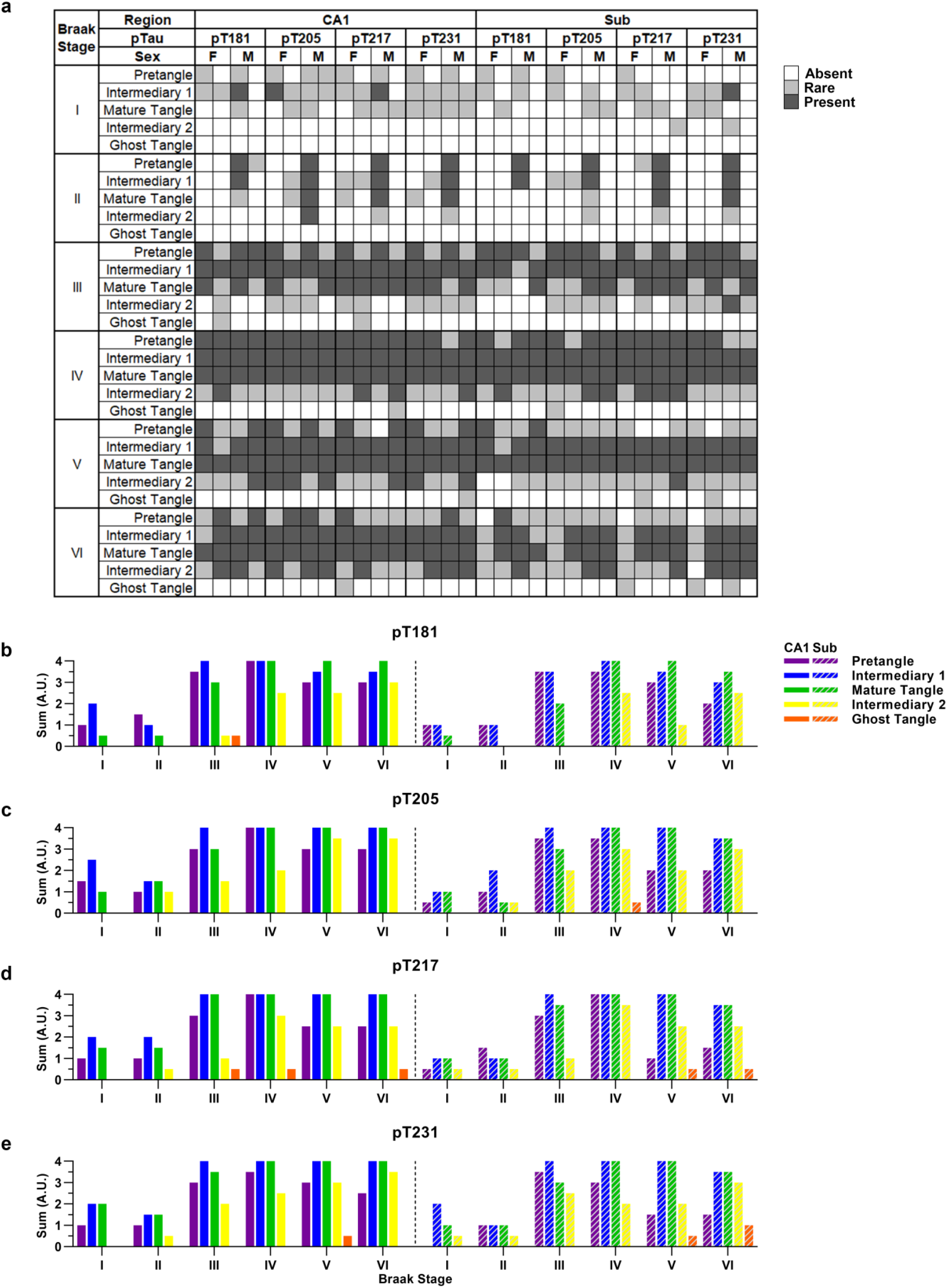
Semi-quantification of neurofibrillary tangle maturity levels for each phosphorylated tau site. **a** Heatmap visualizing semi-quantification from each case in the CA1 and subiculum. Observations are provided for each case stratified by Braak stage (major row) and phosphorylated tau site (major column). Sum of semi-quantification (y-axis) plotted for the CA1 (left) and subiculum (right) for the **b** pT181, **c** pT205, **d** pT217, and **e** pT231 stained slides grouped by Braak stage (x-axis). Acronyms: F, female; M, male; Sub, subiculum; I1, intermediary 1; I2, intermediary 2; A.U.; arbitrary unit.

### Regional tau burden in CA1 and subiculum

We sought to determine the regional differences of tau pathology recognized by these phosphorylated tau antibodies between the CA1 and subiculum. In the past, we observed nearly twice the number of neurofibrillary tangles in the subiculum compared to CA1 in the typical AD subtype, as visualized by thioflavin-S [43]. While tau burden in the subiculum compared to the CA1 was higher for all phosphorylated tau sites, this increase was attenuated compared to the thioflavin-S tangle counts (ratio of subiculum:CA1 for pT181=1.2 [R^2^=0.896]; pT205=1.1, [R^2^=0.932]; pT217=1.2 [R^2^=0.940]; pT231=1.2 [R^2^=0.901]; thioflavin-S=1.7 [R^2^=0.928]) (**Fig. 4**). In the CA1 and subiculum, we found no qualitative difference between the neurofibrillary tangle maturity levels visualized by each phosphorylated tau site. Notably, the frequency of pretangles in Braak stage V/VI were rarer in subiculum compared to the CA1 (**Fig. 3**).

**Figure 4.**
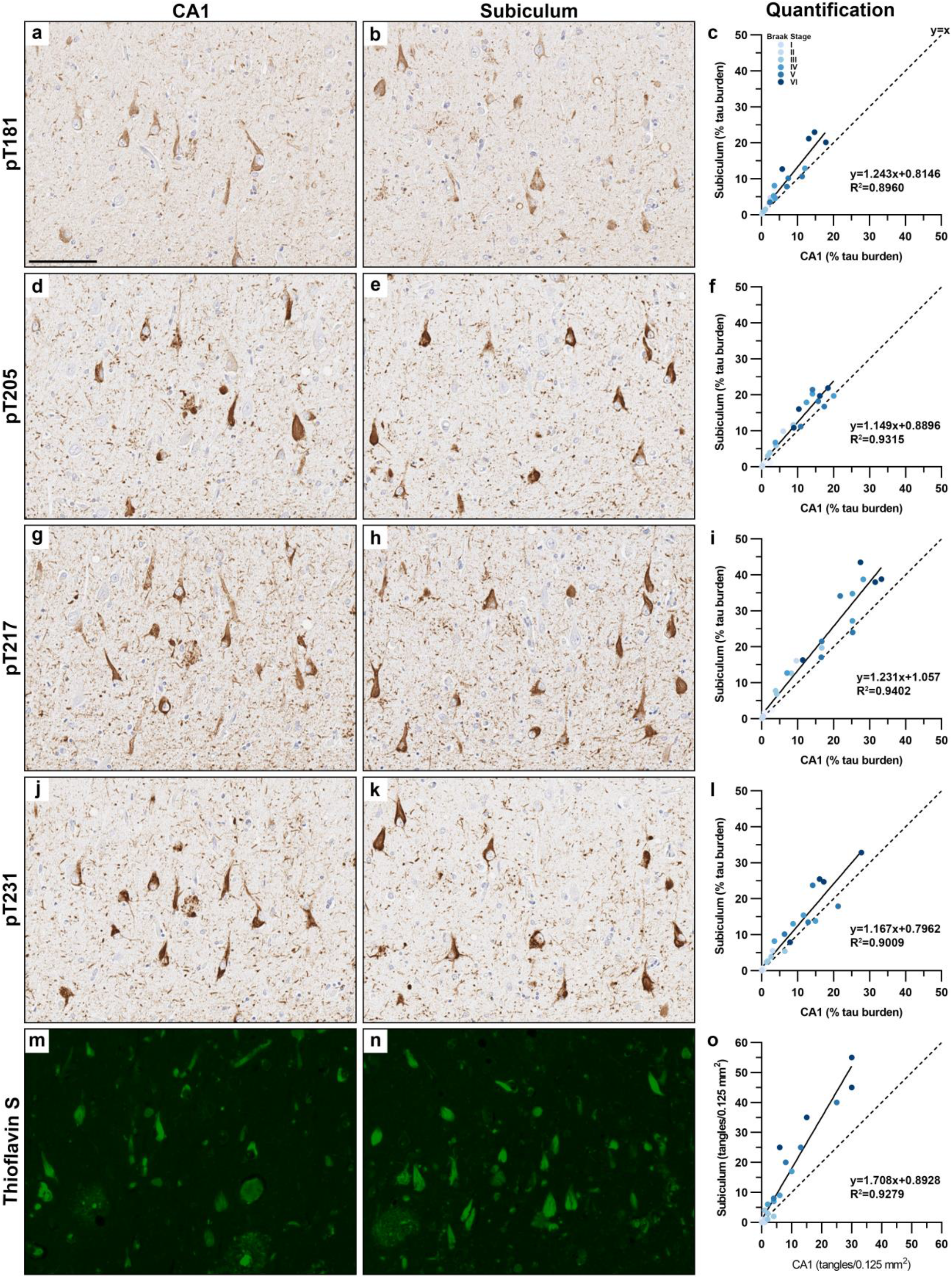
Neurofibrillary tangle pathology in the CA1 and subiculum. Representative images from pT181 in the **a** CA1 and **b** subiculum with **c** quantification. Representative images from pT205 in the **d** CA1 and **e** subiculum with **f** quantification. Representative images from pT217 in the **g** CA1 and **h** subiculum with **i** quantification. Representative images from pT231 in the **j** CA1 and **k** subiculum with **l** quantification. Representative images from Thioflavin S in the **m** CA1 and **n** subiculum with **o** quantification. Tau burden measurements were plotted to evaluate severity differences between the CA1 and subiculum, where the slope of the line (m) provides the subiculum:CA1 ratio from the slope intercept form of a straight line (y=mx+b). A slope of 1 would indicate even distribution (dotted line) and a slope of 2 would be indicative of twice the amount of pathology in subiculum compared to CA1. Tau burden in the subiculum was higher than CA1 for each of the phosphorylated tau sites examined, but differences did not reach the nearly doubled tangle density observed on thioflavin-S. Images are from case 18. Dotted line represents the line of equality (y=x). Scale bar measures 100 µm.

### Global hippocampal evaluation of tau burden

We next sought to determine differences between the phosphorylated tau sites when using the global tau measure, Braak stage [12, 13], to stratify cases. In Braak stages I and II, there were primarily rare pretangles and intermediary 1s (**Fig. 3**). By Braak stage III, there were frequent pretangles, intermediary 1s, and mature tangles, with rare intermediary 2s (**Fig. 3**). From Braak stages IV to VI, there were frequent pretangles, intermediary 1s, mature tangles, and an increasing amount of intermediary 2s; however, there were relatively fewer pretangles in Braak stages V and VI (**Fig. 3**). Ghost tangles were absent to rare across all Braak stages.

In general, tau burden measurements were higher through each Braak stage (**Fig. 5, 6**). pT181 burden increased steadily between Braak stage III and Braak stage VI. pT205 burden increased from Braak stage II to VI, where burden appeared to level off between Braak stages IV and VI. pT217 burden increased from Braak stage II to III, sharply increased from Braak stages III to IV, leveled off from Braak stage IV to V, before increasing again to stage VI. Finally, pT231 burden increased more steadily between Braak stage III to VI.

**Figure 5.**
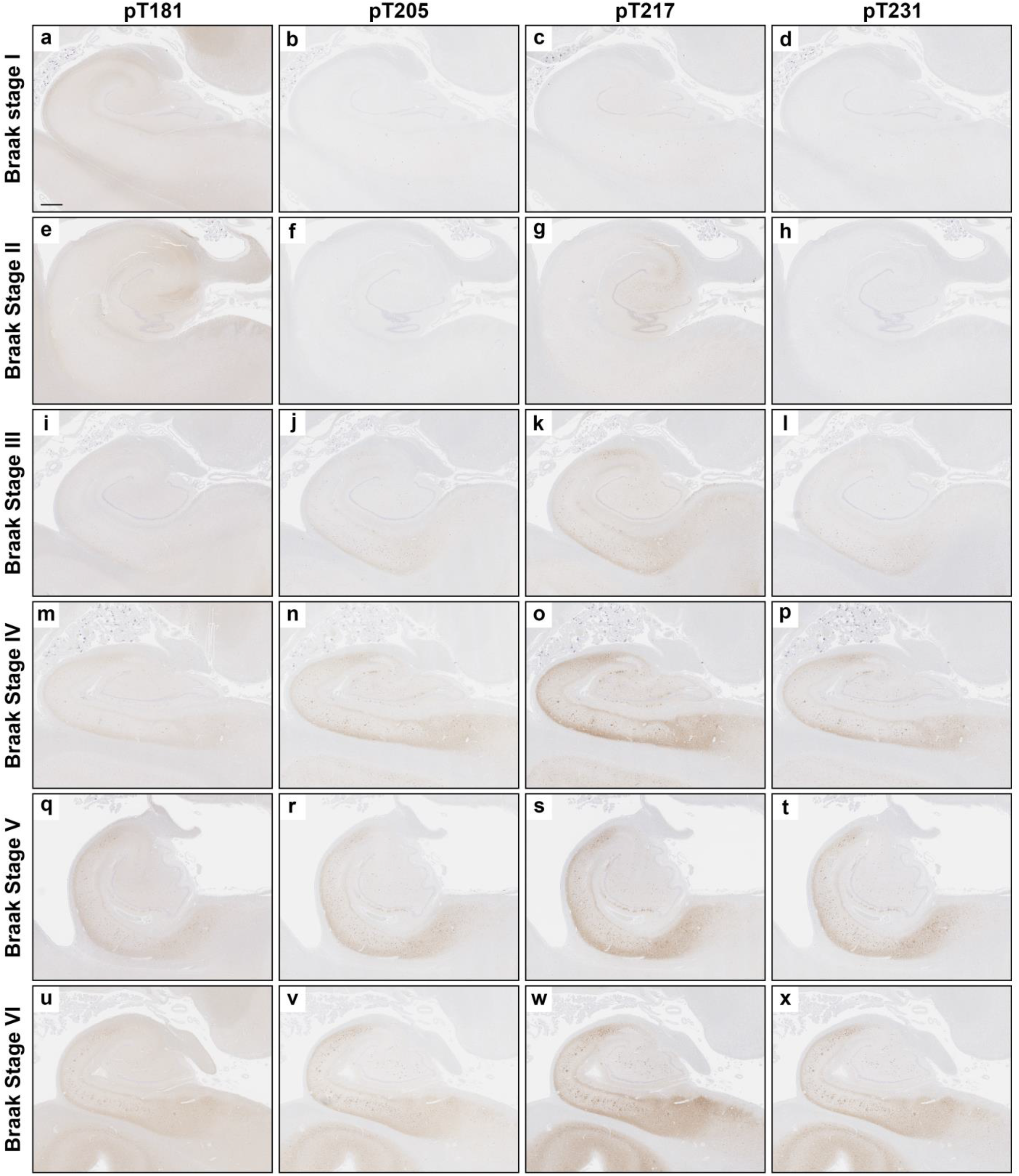
Phosphorylated tau expression generally increases through each Braak Stage. Representative images of immunohistochemistry of the whole posterior hippocampus from **a-d** Braak stage I (case 1), **e-h** Braak stage II (case 5), **i-l** Braak stage III (case 12), **m-p** Braak stage IV (case 13), **q-t** Braak stage V (case 19), and **u-x** Braak stage VI (case 22). Serial sections were stained for **a, e, i, m, q**, and **u** pT181; **b, f, j** ,**n** ,**r**, and **v** pT205; **c, g, k, o, s**, and **w** pT217; and **d, h, l, p, t**, and **x** pT231. As expected, with each increasing Braak stage the hippocampal size decreased. Immunohistochemical staining of tau was visibly increased with each increasing Braak stage.

**Figure 6.**
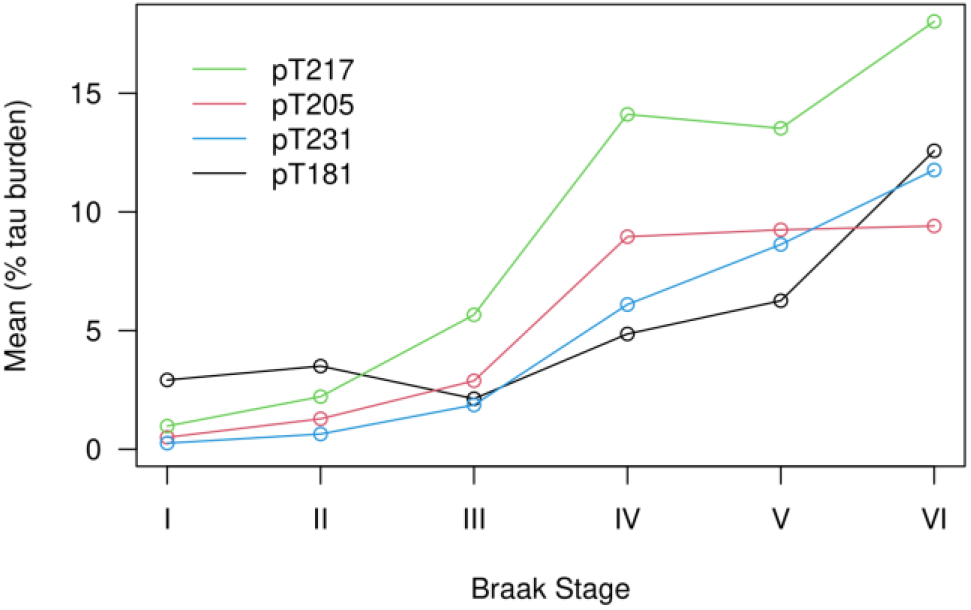
Phosphorylated tau increases through each Braak stage. When comparing tau burden measured across cases at each Braak stages, pT181 burden increased steadily between Braak stage III and Braak stage VI. pT205 burden increased from Braak stage II to VI and appeared to level off between Braak stages IV and VI. pT217 burden increased from Braak stage II to III, and then sharply increased from Braak stages III to IV. pT217 burden leveled off from Braak stage IV to V, before increasing again in stage VI. Finally, pT231 burden increased more steadily between Braak stage III to VI. Line graphs plotted the fitted means using positive pixel count burden for each Braak stage.

## Discussion

Our findings provide neurobiologic evidence that fluid-based phosphorylated tau biomarker sites in the proline-rich domain are present during earlier neurofibrillary tangle maturity. While all neurofibrillary tangle maturity levels were observed in pT181, pT205, pT217, and pT231, there was a predilection toward pretangles, intermediary 1s, and mature tangles. The striking difference in vulnerability between CA1 and subiculum tangle counts observed on thioflavin-S was not as readily observed in tau burden measurements of the phosphorylated tau sites suggesting diminished antibody recognition once tangle pathology matured to advanced levels. Phosphorylated tau burden generally increased as Braak stage increased with some plateauing observed in later Braak stages.

While the temporal sequence of detectable phosphorylated tau levels in fluids is still under investigation [4, 7, 39], evidence suggests that fluid tau levels may elevate earlier than observed uptake on tau positron emission tomography (PET) [23]. Previous tau PET autoradiographic studies suggested that flortaucipir (AV-1451) may recognize middling to advanced neurofibrillary tangle maturity levels [30, 35]. However, there is a lack of in-depth characterization of the neurofibrillary tangle maturity levels [42] recognized by the phosphorylated tau fluid biomarker sites. To fill this knowledge gap, we performed deep phenotyping of neurofibrillary tangle morphology recognized by pT181, pT205, pT217, and pT231 and employed digital pathology to quantify tau burden. We found that all phosphorylated tau sites studied are present primarily in earlier neurofibrillary tangle maturity levels. Their recognition of pretangles, intermediary 1s, and mature tangles may help explain why these biomarker sites are observed earlier during the AD dementia course [4, 8, 21, 37]. To put in context of the current study, and our morphologic characterization across neurofibrillary tangle maturity, we retrospectively examined photomicrographs from previous studies utilizing antibodies that recognized these phosphorylated tau sites. Evaluation of these photomicrographs support our results that pT181 [5, 48], pT205 [50, 51, 54, 60], pT217 [18], and pT231 [5, 10, 22, 32, 34, 58] recognize earlier neurofibrillary tangle maturity levels as we observed labeling of pretangles and mature tangles but not ghost tangles. Based on anticipated projected course in dominantly inherited AD cases, some phosphorylated tau fluid biomarkers were temporally sequenced, suggesting that cerebrospinal fluid levels of p-tau217 may elevate earlier, followed by p-tau181, then p-tau205 [8]. pT231 was recently suggested to be one of the initial phosphorylated sites found in neurons that may develop earlier than pretangles [3], which supports evidence from a plasma study in an autopsy cohort showing plasma p-tau231 levels may distinguish Braak 0 from Braak I-II [4]. Although more work is needed to disentangle the temporal sequence of phosphorylation events, we provide histopathologic evidence that pT181, pT205, pT217, and pT231 are expressed early in the neurofibrillary tangle lifespan. Moreover, accumulation of phosphorylated tau pathology recognizing the proline-rich domain was observed prior to the accumulation of amyloid plaques in three individuals with a Thal amyloid phase 0 (Case #2, #8, #11).

To further investigate differences in regional vulnerability, we elected to examine the CA1 and subiculum, two hippocampal subsectors known to be vulnerable in AD. Based upon observations of nearly twice the number of thioflavin-S positive tangles in the subiculum compared to the CA1 in typical AD [43, 46], our goal was to investigate differences in phosphorylated tau burden in these two regions. However, while phosphorylated tau burden was increased in subiculum compared to CA1, this effect was attenuated compared to thioflavin-S positive tangle counts. We speculate this attenuation may be due to the recognition of pretangles and intermediate 1s by phosphorylated tau sites, which may decrease in number as the disease continues and tau fibrils form into β-pleated sheet structures readily recognized by thioflavin-S. The current study employed digital pathology to quantify phosphorylated tau burden. Thus, we cannot rule out the contribution of neuritic pathology measured by burden analyses not captured by tangle counts performed on thioflavin-S. It is unclear why the subiculum has 1.5-2 times the amount of neurofibrillary tangle burden compared to the CA1 [43, 46]. As the subiculum is the major outflow pathway of the hippocampus that receives multiple connections from CA1 and entorhinal cortex [36], these anatomical differences may confer heightened vulnerability to advanced neurofibrillary tangle maturity. Evaluation in the context of reciprocal connections [29] or perhaps populations of excitatory neurons [27] may warrant future investigation.

We further investigated hippocampal differences in these phosphorylated tau sites across a global measure of tau distribution, Braak stage [12, 13]. As expected in Braak stages I-II, we found minimal neurofibrillary tangle pathology primarily consisting of pretangles, intermediary 1s, and rarely mature tangles. However, by Braak stage III and onwards, there was an increase in the neurofibrillary tangle pathology with an increase in mature tangle and rare observations of intermediary 2. By Braak stage V, there was a notable decrease in pretangle population. This is not unexpected as the number of unaffected neurons with potential to accumulate pathology decreases as pathology burden increases [16]. Ghost tangles were not commonly observed, which may indicate the loss of the phosphate group at the advanced maturity levels. We hypothesize this is because as ghost tangles are the remnants of mature tangles once the neuron has died [2, 49] they are no longer contained by a membrane and as such are exposed to the surrounding environment including phosphatases. In a neuropathologically diagnosed cohort, plasma p-tau181 and p-tau231 levels were found to increase with Braak stage [4] and suggested to plateau in very advanced stages of disease [26]. Our findings provide support of the observed plateauing, suggesting that as the shift from early to advanced tangle maturity occurs recognition by these phosphorylated tau sites in the proline-rich domain may be diminished.

Our study is not without limitations. One such is sample size; while n=4 cases per Braak stage is adequate for the morphology characterization and semiquantitative methods of neurofibrillary tangle maturity, larger sample sizes are necessary to evaluate and interpret tau burden differences between Braak stages. Another limitation concerns the thickness of tissue sections. It is difficult to be certain of intermediary 2 tangles, as it is possible that the nucleus is out of plane of section. To overcome this, thicker sections can be used to include the entire neurofibrillary tangle. In the current study, we observed lighter staining by the pT181 antibody compared to the other phosphorylated tau sites. As antibody affinity may affect immunohistochemical burden analyses, caution is warranted in interpretation of which phosphorylated tau site is increased the earliest in the current study. Additionally, we chose to focus on the hippocampus as this region is well-characterized in terms of neurofibrillary tangle maturity [6, 42, 55]. As neurofibrillary tangle maturity is not well-characterized in cortical regions, future studies should consider characterizing neurofibrillary tangle maturity levels in these regions to facilitate investigation of these phosphorylated tau fluid biomarker sites.

In conclusion, we performed a deep postmortem characterization of four phosphorylated tau fluid biomarker sites (pT181, pT205, pT217, and pT231) that are reliably elevated in AD dementia. To our knowledge, this is the first time these four phosphorylated tau sites are characterized together in the postmortem brain to evaluate recognition of neurofibrillary tangle maturity. While all neurofibrillary tangle maturity levels were visualized by each of the four phosphorylated tau sites, we observed a predilection towards earlier neurofibrillary tangle maturity with extremely rare observations of ghost tangles. Although fluid biomarkers do not provide regional involvement captured by tau PET imaging, our study demonstrates these phosphorylated tau sites in the proline-rich domain readily recognize early aspects of tangle maturity and may provide keen insight into the initial phase of neurofibrillary tangle changes.

## Acknowledgements

We are grateful to Virginia Phillips, Ariston Librero, Jo Landino, Jessica Tranovich, Janisse Cabrera, and the Cytometry and Imaging Lab for histologic and imaging support.

## Compliance with Ethical Standards

### Funding

The investigators are supported by grants from National Institute on Aging (R01 AG054449, P30 AG062677, RF1 AG069052, U01 AG057195, and P50 AG047266) and the Florida Department of Health, Ed and Ethel Moore Alzheimer’s Disease Research Program (8AZ06, 20A22).

### Declarations

All brains were acquired with appropriate ethical approval, and the research performed on postmortem samples was approved by the Mayo Clinic Research Executive Committee.

#### Abbreviations

AD: Alzheimer’s disease
ARTAG: Aging-related tau astrogliopathy
CA: Cornu ammonis
FLAME: Florida Autopsied Multi-Ethnic
I1: Intermediary 1
I2: Intermediary 2
p: Phosphorylation
PA: Pathological aging
PART: Primary age related tauopathy
PET: Positron emission tomography
SC: Senile change
T: Threonine
TSA: Thorn shaped astrocytes
VaD: Vascular disease

## Supplementary Information

**Supplementary Table 1.**
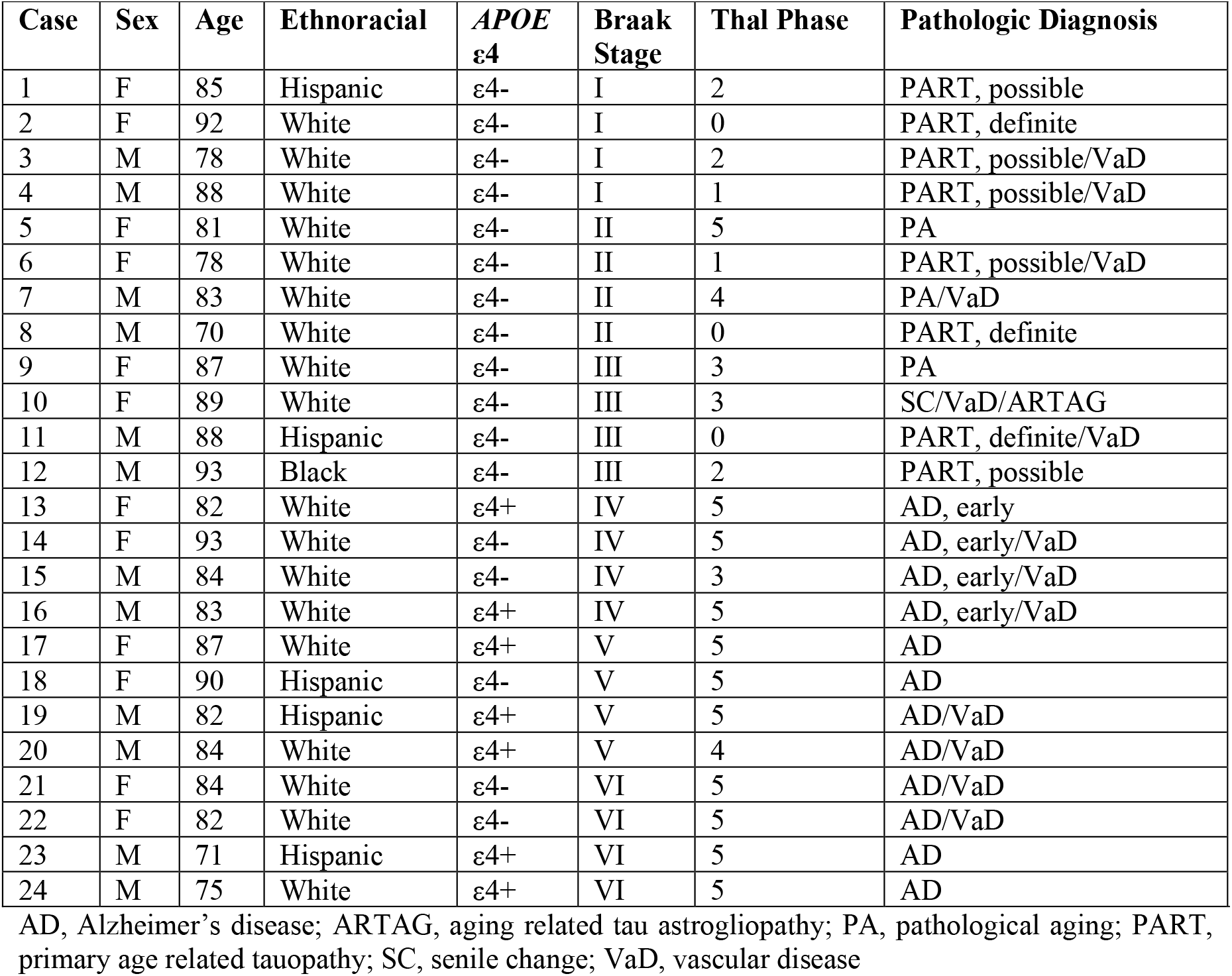
Case demographics

| Case | Sex | Age | Ethnoracial | APOE<br>ε4 | Braak<br>Stage | Thal Phase | Pathologic Diagnosis |
| --- | --- | --- | --- | --- | --- | --- | --- |
| 1 | F | 85 | Hispanic | ε4- | I | 2 | PART, possible |
| 2 | F | 92 | White | ε4- | I | 0 | PART, definite |
| 3 | M | 78 | White | ε4- | I | 2 | PART, possible/VaD |
| 4 | M | 88 | White | ε4- | I | 1 | PART, possible/VaD |
| 5 | F | 81 | White | ε4- | II | 5 | PA |
| 6 | F | 78 | White | ε4- | II | 1 | PART, possible/VaD |
| 7 | M | 83 | White | ε4- | II | 4 | PA/VaD |
| 8 | M | 70 | White | ε4- | II | 0 | PART, definite |
| 9 | F | 87 | White | ε4- | III | 3 | PA |
| 10 | F | 89 | White | ε4- | III | 3 | SC/VaD/ARTAG |
| 11 | M | 88 | Hispanic | ε4- | III | 0 | PART, definite/VaD |
| 12 | M | 93 | Black | ε4- | III | 2 | PART, possible |
| 13 | F | 82 | White | ε4+ | IV | 5 | AD, early |
| 14 | F | 93 | White | ε4- | IV | 5 | AD, early/VaD |
| 15 | M | 84 | White | ε4- | IV | 3 | AD, early/VaD |
| 16 | M | 83 | White | ε4+ | IV | 5 | AD, early/VaD |
| 17 | F | 87 | White | ε4+ | V | 5 | AD |
| 18 | F | 90 | Hispanic | ε4- | V | 5 | AD |
| 19 | M | 82 | Hispanic | ε4+ | V | 5 | AD/VaD |
| 20 | M | 84 | White | ε4+ | V | 4 | AD/VaD |
| 21 | F | 84 | White | ε4- | VI | 5 | AD/VaD |
| 22 | F | 82 | White | ε4- | VI | 5 | AD/VaD |
| 23 | M | 71 | Hispanic | ε4+ | VI | 5 | AD |
| 24 | M | 75 | White | ε4+ | VI | 5 | AD |

AD, Alzheimer's disease; ARTAG, aging related tau astrogliopathy; PA, pathological aging; PART, primary age related tauopathy; SC, senile change; VaD, vascular disease

**Supplementary Table 2.** Positive pixel count macro specifications for pT181, pT205, pT217, and pT231 stained sections.

| Algorithm | Positive Pixel Count |
| --- | --- |
| Version | 9.1 |
| View Width | 1000 |
| View Height | 1000 |
| Overlap Size | 0 |
| Image Zoom | 1. |
| Classifier | None |
| Class List |  |
| Classifier Neighborhood | 0 |
| Pixel Area (millimeter-squared) | 2.52506e-007 |
| Hue Value (Center) | 0.1 |
| Hue Width | 0.1 |
| Color Saturation Threshold | 8.e-002 |
| Intensity Threshold WEAK (Upper Limit) | 235 |
| Intensity Threshold WEAK (Lower Limit) | 235 |
| Intensity Threshold MEDIUM (Upper Limit) | 235 |
| Intensity Threshold MEDIUM (Lower Limit) | 235 |
| Intensity Threshold STRONG (Upper Limit) | 235 |
| Intensity Threshold STRONG (Lower Limit) | 0 |
| Intensity Threshold Negative Pixels | -1 |

**Supplementary Table 3.**
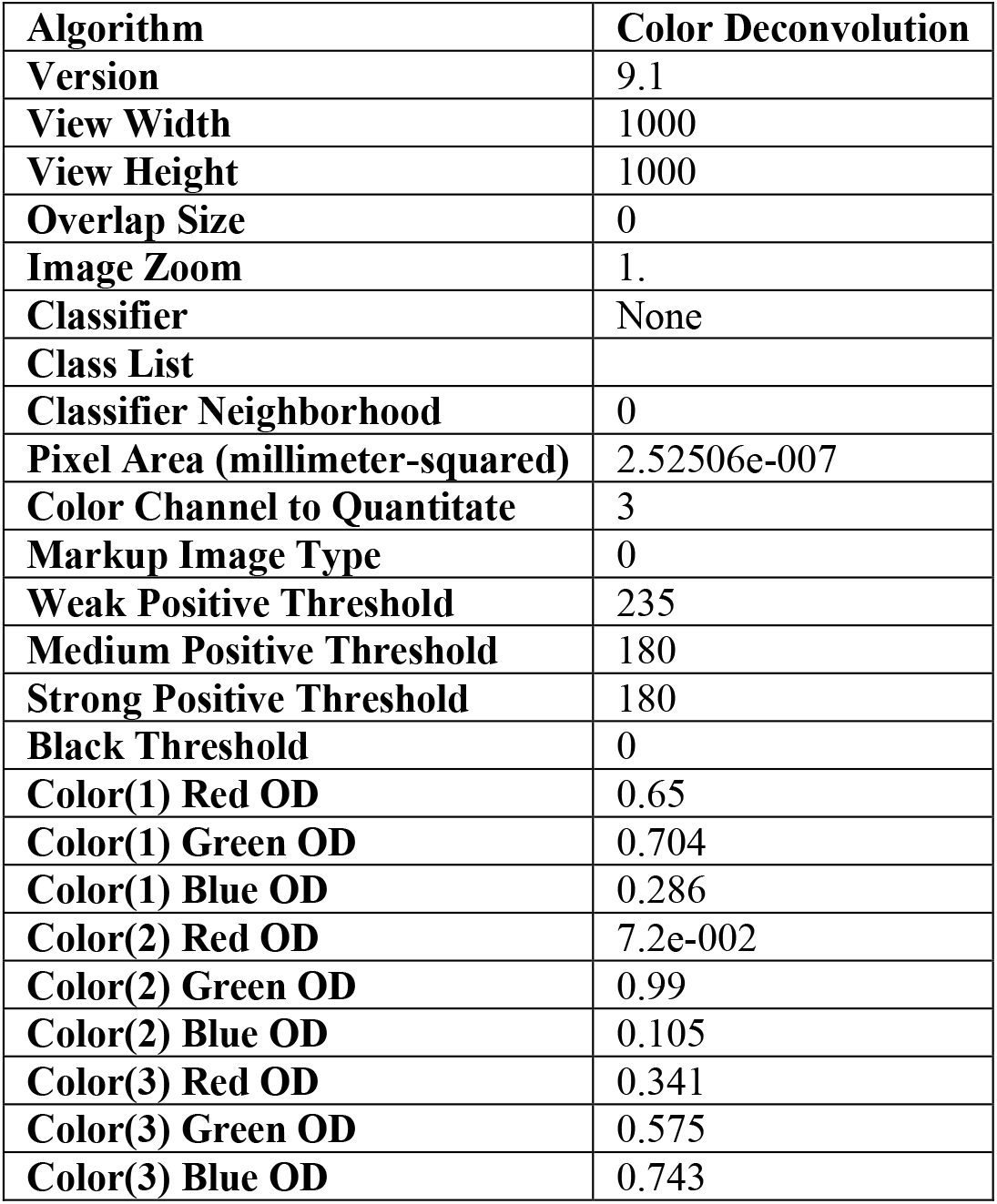
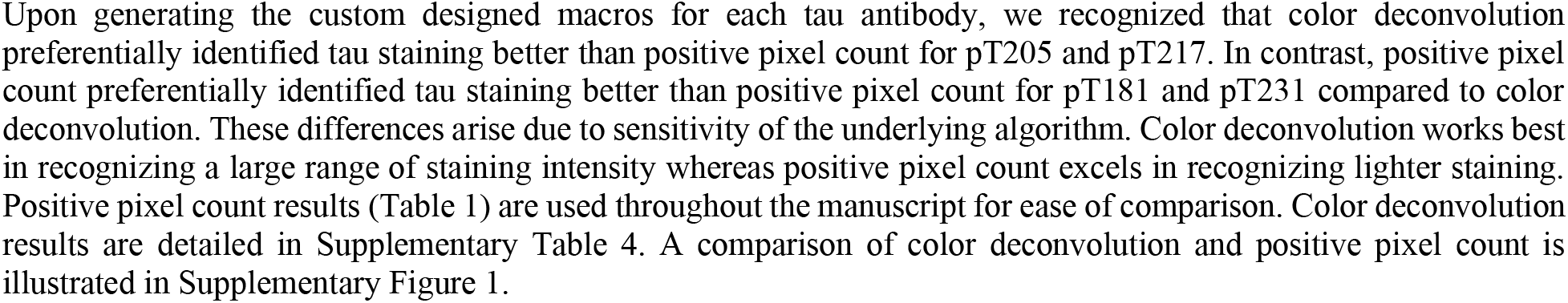
Color deconvolution macro specifications for pT181, pT205, pT217, and pT231 stained sections.

**Supplementary Table 4.** Positive pixel count burden.

| Characteristic | Braak I | Braak II | Braak III | Braak IV | Braak V | Braak VI | p-value |
| --- | --- | --- | --- | --- | --- | --- | --- |
| Whole hippocampus measures |  |  |  |  |  |  |  |
| Color Deconvolution |  |  |  |  |  |  |  |
| pT181 burden, % | 0.58 (0.27, 4.6) | 1.3 (0.21, 3.9) | 0.94 (0.43, 1.9) | 1.8 (1.4, 4.7) | 4.2 (0.77, 5.2) | 6.1 (5.7, 6.8) | <0.001 |
| pT205 burden, % | 0.31 (0.12, 0.48) | 0.12 (0.031, 3.1) | 1.7 (0.83, 3.6) | 5.6 (1.6, 7.3) | 5.7 (4.7, 7.3) | 6.2 (4.1, 7.2) | <0.001 |
| pT217 burden, % | 0.51 (0.26, 1.2) | 0.41 (0.049, 5.0) | 3.4 (1.9, 7.0) | 11 (3.3, 12) | 9.2 (7.4, 12) | 14 (6.2, 17) | <0.001 |
| pT231 burden, % | 0.17 (0.10, 0.32) | 0.079 (0.024, 1.8) | 1.2 (0.63, 2.4) | 4.5 (1.8, 5.8) | 6.1 (3.0, 9.3) | 8.3 (4.6, 11) | <0.001 |
| CA1 measures |  |  |  |  |  |  |  |
| Color Deconvolution |  |  |  |  |  |  |  |
| pT181 burden, % | 0.39 (0.12, 1.7) | 0.73 (0.091, 1.5) | 0.94 (0.38, 1.9) | 1.9 (1.5, 6.9) | 4.6 (1.1, 6.7) | 7.1 (2.6, 10) | <0.001 |
| pT205 burden, % | 0.38 (0.19, 1.3) | 0.14 (0.031, 4.2) | 2.3 (1.3, 5.7) | 7.6 (2.2, 11) | 9.0 (7.6, 11) | 8.9 (5.2, 12) | <0.001 |
| pT217 burden, % | 0.48 (0.39, 2.1) | 0.25 (0.044, 6.4) | 4.3 (2.7, 12) | 18 (4.1, 19) | 14 (11, 17) | 21 (7.3, 24) | <0.001 |
| pT231 burden, % | 0.18 (0.16, 0.61) | 0.082 (0.023, 2.3) | 1.5 (1.1, 4.4) | 6.8 (2.0, 10) | 9.5 (3.9, 15) | 11 (4.9, 18) | <0.001 |
| Subiculum measures |  |  |  |  |  |  |  |
| Color Deconvolution |  |  |  |  |  |  |  |
| pT181 burden, % | 0.47 (0.21, 3.1) | 1.1 (0.12, 2.2) | 1.2 (0.56, 2.2) | 3.0 (2.4, 6.9) | 5.5 (1.4, 7.3) | 11 (6.6, 13) | <0.001 |
| pT205 burden, % | 0.63 (0.17, 0.73) | 0.16 (0.037, 6.3) | 3.4 (2.0, 6.8) | 10 (3.9, 11) | 10 (8.1, 15) | 12 (7.0, 15) | <0.001 |
| pT217 burden, % | 0.96 (0.42, 1.7) | 0.41 (0.056, 11) | 7.2 (4.8, 13) | 21 (7.6, 25) | 15 (13, 25) | 29 (12, 33) | <0.001 |
| pT231 burden, % | 0.34 (0.20, 0.54) | 0.098 (0.028, 4.2) | 2.4 (1.8, 3.6) | 9.5 (5.1, 11) | 11 (6.0, 17) | 17 (5.9, 22) | <0.001 |

**Supplementary Figure 1.**
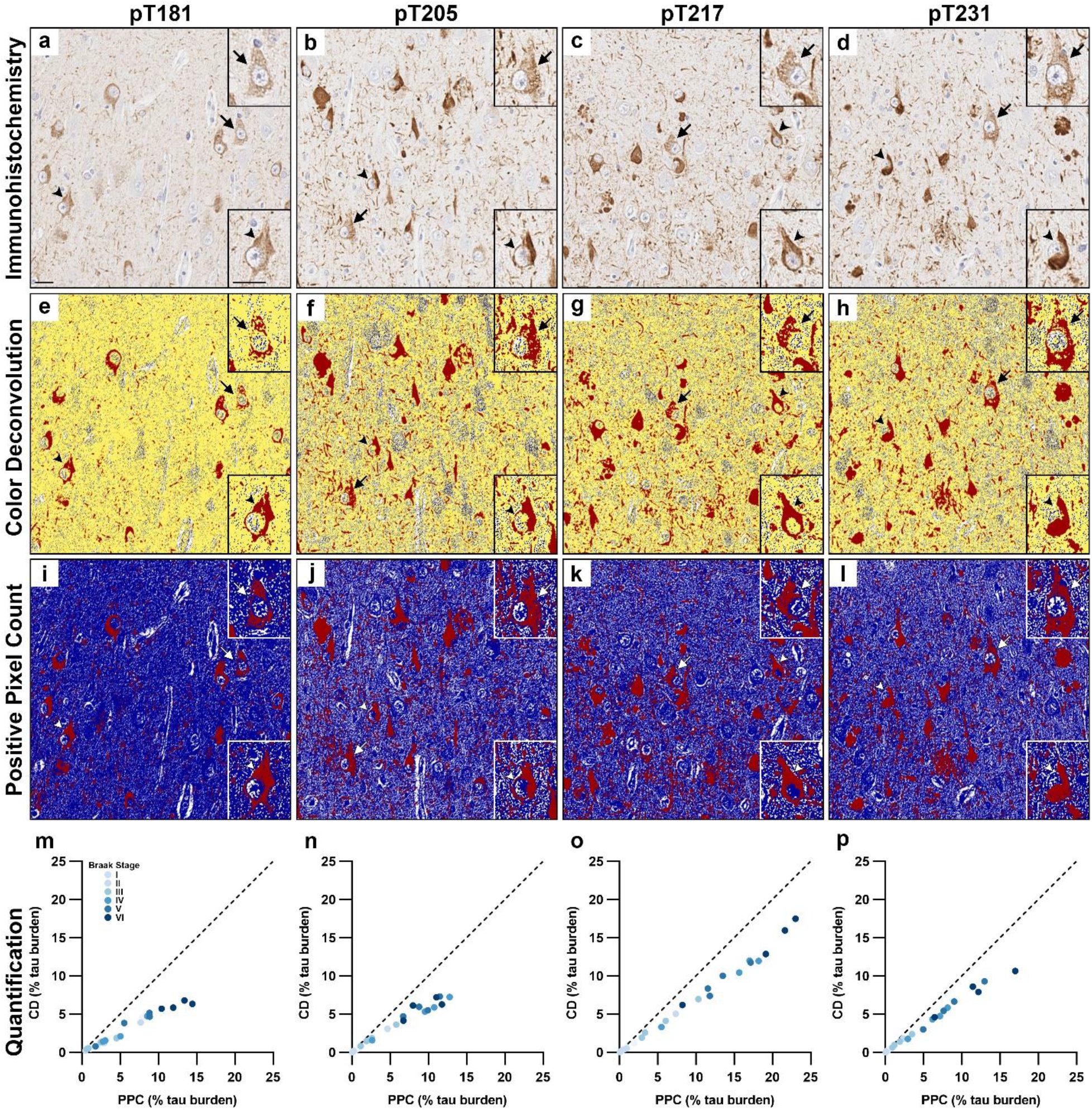
Comparison of color deconvolution and positive pixel count. Immunohistochemistry of the CA1 region from case 20 stained for **a** pT181, **b** pT205, **c** pT217, and **d** pT231. Color deconvolution macro overlay for **e** pT181, **f** pT205, **g** pT217, and **h** pT231. Positive pixel count macro overlay for **i** pT181, **j** pT205, **k** pT217, and **l** pT231. Comparison of color deconvolution and positive pixel count macros for **m** pT181, **n** pT205, **o** pT217, and **p** pT231. Images are from case 20. Dotted line represents the line of equality (y=x). Arrow, earlier tangle maturity level (i.e. pretangle or intermediary 1); arrowhead, mature tangle. Scale bar measures 25 µm.

**Supplementary Figure 2.**
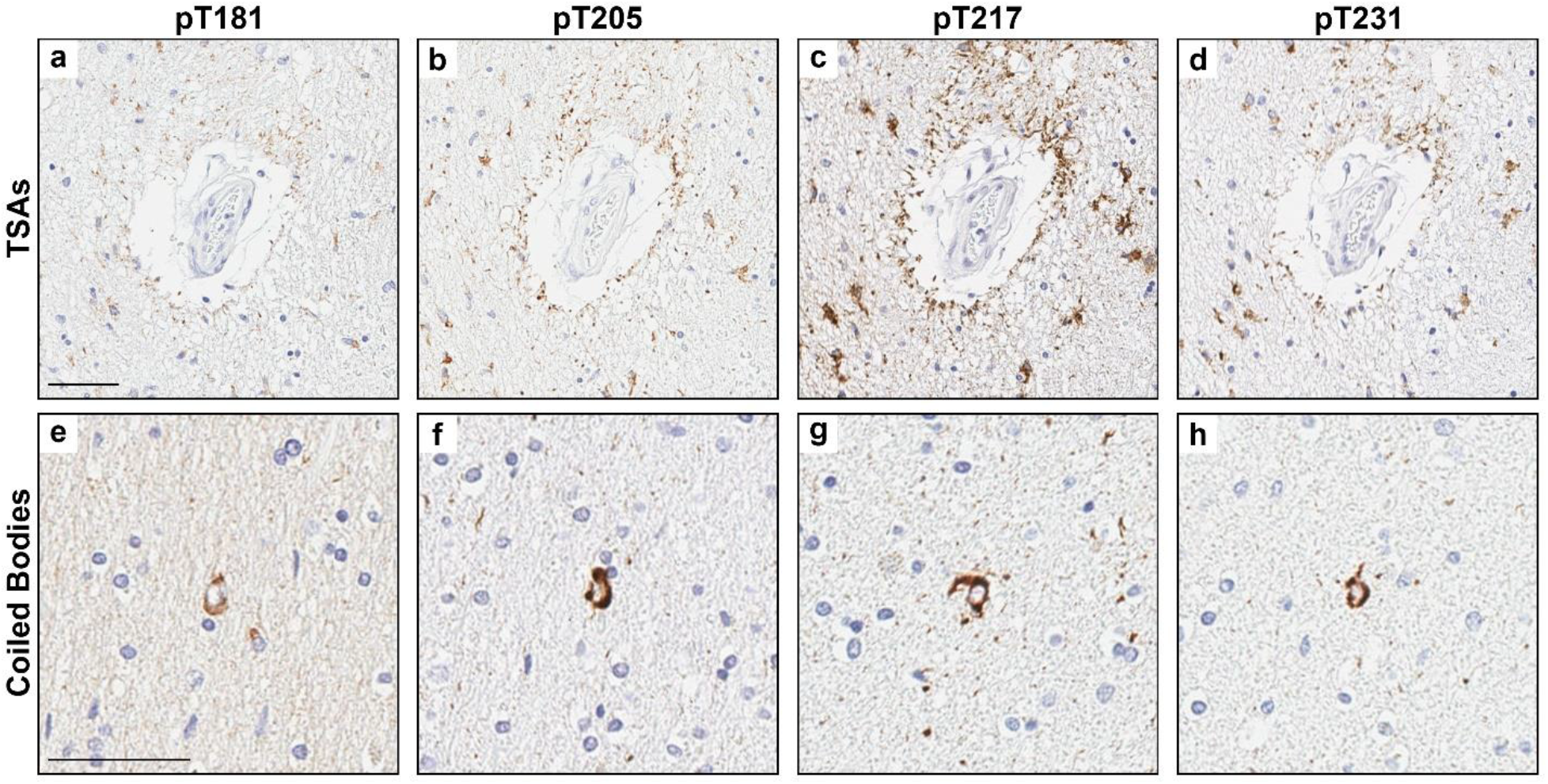
Examples of additional tau pathology observed with each phosphorylated tau site. **a-d** Thorn shaped astrocytes (TSAs), a feature of ARTAG (case 12) and **e-h** Coiled bodies, a feature of grains disease (case 18). **a, e** were stained with the pT181 antibody. **b, f** were stained with the pT205 antibody. **c, g** were stained with the pT217 antibody. **d, h** were stained with the pT231 antibody. Scale bar measures 50 µm.

